# Efect of Sec61 interaction with Mpd1 on Endoplasmic Reticulum- Associated Degradation

**DOI:** 10.1101/411173

**Authors:** Fábio Pereira, Mandy Rettel, Frank Stein, Mikhail M. Savitski, Ian Collinson, Karin Römisch

## Abstract

Proteins that misfold in the endoplasmic reticulum (ER) are transported back to the cytosol for ER-associated degradation (ERAD). The Sec61 channel is one of the candidates for the retrograde transport conduit. Channel opening from the ER lumen must be triggered by ERAD factors and substrates. Here we identified new lumenal interaction partners of Sec61 by chemical crosslinking and mass spectrometry. In addition to known Sec61 interactors we detected ERAD factors including Cue1, Ubc6, Ubc7, Asi3, and Mpd1. We show that the CPY* ERAD factor Mpd1 binds to the lumenal Sec61 hinge region. Deletion of the Mpd1 binding site reduced the interaction between both proteins and caused an ERAD defect specific for CPY* without affecting protein import into the ER or ERAD of other substrates. Our data suggest that Mpd1 binding to Sec61 is a prerequisite for CPY* ERAD and confirm a role of Sec61 in ERAD of misfolded secretory proteins.

## Introduction

In eukaryotes about 30% of all proteins constitute secretory pathway cargo (Ghaemmaghami et al., 2003). These proteins are transported into the ER by the conserved heterotrimeric Sec61 channel formed by Sec61, Sbh1, and Sss1 in yeast (Sec61*α*, Sec61*β*, Sec61*γ* in mammals) (Johnson and van Waes, 1999). During targeting and translocation the Sec61 channel interacts with multiple other protein complexes on its cytosolic face and in the ER membrane such as the ribosome, the SRP receptor, the Sec63 complex, oligosaccharyl transferase, and signal peptidase (Kalies et al., 1994; Brodsky et al., 1995; Jadhav et al., 2015; Scheper et al., 2003; Kalies et al., 1998). If proteins fail to fold in the ER, they trigger the Unfolded Protein Response (UPR), unless they are transported back to the cytosol for ERAD (Pilla et al., 2017; Römisch, 2017). Although this process has been intensely studied for over 20 years, the identity of the retrograde transport channel is still controversial. The first and most investigated candidate is the Sec61 channel (Römisch, 2017). The E3 ubiquitin ligase Hrd1 and the pseudorhomboid proteases Der1 and Dfm1 have been proposed more recently as ERAD channels (Mehnert et al., 2014; Neal et al., 2018). The Sec61 channel has been shown to interact with Hrd1, and Hrd1 with Der1, so these proteins may also operate together in transporting ERAD substrates to the cytosol (Ng et al., 2007; Carvalho et al., 2006; Römisch, 2017). If the Sec61 channel were involved in retrograde transport of ERAD substrates, it would have to interact with ERAD factors targeting ERAD substrates to its lumenal end. While Sec61 interaction with ERAD substrates has been shown (Pilon et al., 1997; Schäfer and Wolf, 2009), the only known ER lumenal ERAD factor that is known to interact with Sec61 is the Hsp70 BiP (Schäuble et al., 2012). Here we have used chemical crosslinking and mass spectrometry to identify new interactors of Sec61 with specific focus on ERAD-relevant and lumenal interactors in order to better understand the role of the Sec61 channel in this process.

## Results and Discussion

To identify new lumenal interaction partners of Sec61 we used a functional *sec61* mutant with a unique cysteine in its large lumenal loop 7 (Fig. 1A) (Kaiser and Römisch, 2015). Using hetero- bifunctional non-cleavable (SMPH) or cleavable (LC-SPDP) crosslinkers with a cysteine-reactive group and a NH_2_-reactive group to crosslink yeast microsomes, as described in Materials & Methods, we found additional bands in the crosslinking patterns to Sec61S353C compared to wildtype Sec61-suggesting bound lumenal interactors (Fig. 1B, C, arrows). Amongst those was Sec63, a well-characterized J-domain protein that contributes to both co- and posttranslational import into the ER and to ERAD (Fig. 1B) (Brodsky et al., 1995; Servas and Römisch, 2013). While pretreatment of microsomes with urea had no effect on the Sec61S353C-associated proteins (Fig. 1D, lanes 4-6), extraction of microsomes with sodium carbonate resulted in reduced crosslinking to Sss1 which is known to be partially carbonate-extractable (Esnault et al., 1994) and to Sec63 (Fig. 1D, lanes 10-12). Our data suggest an interaction between the Sec63 lumenal J-domain or N-terminus with Sec61 loop7.

For enrichment of Sec61-crosslinked proteins we tagged the N-termini of Sec61 and Sec61S353C with 14-His which had no effects on growth, expression levels, or tunicamycin-sensitivity and UPR induction (not shown), indicating no perturbance of ER proteostasis. Crosslinking patterns were not affected by the tagging (Fig. 1E). Sec61- and Sec61S353C-crosslinked proteins were purified from 500 eq lysed microsomes on a nickel column and eluted with imidazole (Fig. 2A). Fractions 3-10 of the eluates were pooled and proteins analyzed by mass spectrometry. Proteins were accepted as interactors if there was at least a 3-fold enrichment compared to the uncrosslinked sample (Fig. 2B). In total, we identified 353 proteins that were copurifying with Sec61 in the crosslinked samples (supplementary table to Fig. 2). While the enrichment pattern was sample- and crosslinker-dependent (supplementary table to Fig. 2), the absolute abundance of proteins in the ER did not affect interaction with Sec61 (Fig. 2C) suggesting that the interactions we detected were specific. We detected all subunits of the Sec complex in the ER membrane, SRP receptor, Snd3, and several subunits of oligosaccharyl transferase (supplementary table to Fig. 2) (Aviram et al., 2016; Wang and Dobberstein, 1999). In the same significance range we found a number of new interaction partners of Sec61 that were ERAD relevant: Asi3, Ubc6, Ubc7, Cue1, Ubx7, Ubp1, Rpt2, ER-membrane complex (EMC) subunits, and Mpd1, suggesting close physical contact of the Sec61 channel with ERAD machinery (Foresti et al., 2014; Römisch, 2005; Vembar and Brodsky, 2008; Ng et al., 2007; Baker et al., 1992; Christianson et al., 2011; Grubb et al., 2012).

We then decided to investigate the interaction of Sec61 with Mpd1, a known ERAD factor of the well-characterized ERAD substrate CPY* (Grubb et al., 2012). Our xQuest/xProphet analysis of crosslinked peptides suggested a direct interaction of Mpd1 C59 with K209 in lumenal loop5 of Sec61 which constitutes the hinge region around which the N-terminal half of Sec61 swings during channel opening (Fig. 3A, upper) (Leitner et al., 2014; Voorhees and Hegde, 2016). Comparison of Sec61 loop5 with SecY loop5 of bacteria and archaea revealed a substantial extension of loop5 in eukaryotes including the crosslinking site of Mpd1 (Fig. 3A, middle & lower). We hypothesized that the eukaryotic extensions in loop5 might serve as docking sites for ERAD factors to facilitate opening of the Sec61 channel from the lumen for export of ERAD substrates. To test this hypothesis we deleted sections of the Sec61 hinge including the Mpd1 contact site to create a smaller vestigial hinge within Sec61, similar to the SecY counterpart (Fig. 3A, middle & lower), and investigated the effects on protein transport into the ER and ERAD. While deletion1 caused temperature- and cold-sensitivity alone and in combination with deletion2 (Fig. 3B), steady-state expression levels of all hinge mutants were like wildtype (Fig. 4F), and there was no effect on co- or posttranslational protein import into the ER (Fig. 3C).

As only the double mutant *sec61del1/2* showed a moderate tunicamycin-sensitivity (Fig. 3B) and slightly induced UPR (Fig. 3 - Figure supplement 1), ER-proteostasis was not dramatically compromised in the mutants excluding gross ERAD defects. This was confirmed by normal ERAD kinetics for the KHN, KWW, and pΔgp*α*F substrates in the mutants (Fig. 4A,B,C) (Pilon et al., 1997; Vashist and Ng, 2004). CPY* degradation, however, was compromised in *sec61del1* which lacks the contact site for Mpd1 (Fig. 4D, magenta). In contrast, *sec61del2* barely affected CPY* degradation (Fig. 4D, red). The *sec61del1/2* mutant had an intermediate phenotype (Fig. 4D, green) which may suggest that it was not just the absence of specific amino acids deleted in *sec61del1*, but also the distortion of the hinge by the deletion that caused the CPY* ERAD defect (Fig. 3A, lower). In *sec61del1/2* this distortion is partially compensated (Fig 3A, lower).

To directly confirm that the Mpd1 interaction with Sec61 was compromised in the *sec61* hinge mutants we prepared radiolabelled microsomes from wildtype, *sec61S353C* and *sec61* hinge mutant strains expressing HA-tagged Mpd1 and performed sequential immunoprecipitations with Sec61 and HA-antibodies. In all hinge mutants less Mpd1 was associated with Sec61 compared to wildtype or Sec61S353C (Fig. 4E), but it was not possible to correlate that amount of Mpd1 bound to mutant Sec61 with the degree of the CPY* ERAD defect (compare Figs. 4D, 4E). To exclude that the *sec61* hinge mutants reduced biogenesis of the ER ubiquitin ligase Hrd1 and its cofactor Hrd3 we performed quantitative immunoblots for both proteins and found that they were expressed equally in wildtype and mutant cells (Fig. 4F).

Collectively, our data suggest that interaction of the CPY* ERAD factor Mpd1 with the Sec61 hinge region in loop5 contributes to export and degradation of this substrate. Our results are consistent with the view that Sec61 forms part of an export complex in the ER membrane for misfolded protein transport to the cytosol (Fig. 4G). The extended hinge in Sec61 compared to SecY (Fig. 3A) may serve to activate and open the channel from the lumen for intercalation and subsequent transport of CPY* to the cytosol (Fig. 4G).

## Materials and Methods

*S. cerevisiae* strains used in this study are listed in Table 1, plasmids in Table 2, primers in Table 3, and antibodies in Table 4

### Growth of *S. cerevisiae*

*S. cerevisiae* cells were grown at 30°C in YPD or in SC medium with continuous shaking at 220 rpm. Cells on solid medium were also grown at 30°C if not stated otherwise. To test temperature sensitivity, cells were counted and serial dilutions were prepared. A volume of 5 µl of each dilution (containing 10^4^ – 10 cells) was pipetted onto YPD plates. To test tunicamycin (Tm) (SIGMA) sensitivity, cells were grown on YPD plates supplemented with 0, 0.25 or 0.5µg/ml Tm. Plates were incubated at indicated temperatures for 3 days.

### Yeast Microsome Preparation

The isolation of rough microsomal membranes from *S. cerevisiae* was done as in Pilon et al. (1997) and membranes aliquoted at an *OD*_280_=30, snap-frozen in liquid nitrogen, and stored at -80°C. Microsome amounts are referred to as equivalents (eq) in which 1 eq = 1 µl of microsomes at an *OD*_280_ of 50 (Walter et al., 1981).

To prepare radiolabeled ER vesicles, 7 *OD*_600_ of early log-phase cells were incubated in synthetic minimal media supplemented appropriately and lacking methionine, cysteine, and ammonium sulfate for 30 min at 30°C, 220 rpm. Cells were labelled with 6,5 MBq [35S] methionine/cysteine (Express Labeling, PerkinElmer) mix for 30 min. After labelling, cells were immediately washed twice with Tris-Azide Buffer (20 mM Tris-HCl, pH 7.5, 20 mM sodium azide). Cells were then incubated in 100 mM Tris-HCl, pH 9, 10 mM DTT for 10 min at room temperature, sedimented, and resuspended in 300 µl of 2 x JR Lysis Buffer (40 mM Hepes-KOH, pH 7.4, 400 mM sorbitol, 100 mM KOAc, 4 mM EDTA, 1 mM DTT, 1 mM PMSF) (Pilon et al., 1997). Acid-washed glass beads (1/2 volume) were added and the sample submitted at 2 cycles of 1 min bead-beating (Mini-beadbeater-16, BioSpec) with 2 min of incubation on ice after each cycle. From this point on, all samples were kept at 4°C. Beads were washed 3 times with 300 µl of B88, pH 7.2 (20 mM Hepes-KOH pH 6.8, 250 mM sorbitol, 150 mM KOAc, 5 mM Mg(OAc)_2_). Washes were pooled and sedimented for 2 min at 1,500 x g and the microsome-containing supernatant was transferred to a clean tube. Microsomes were then sedimented at 16,000 x g for 10 min, washed and resuspended in 200 µl B88, pH 7.2. Crude radiolabelled ER vesicles were then aliquoted (50 µl), flash frozen in liquid nitrogen, and stored at -80°C.

### Chemical Crosslinking

Microsomes (17 eq) were washed and resuspended in B88 (20 mM Hepes-KOH, 250 mM sorbitol, 150 mM KOAc, 5 mM Mg(OAc)_2_). For SMPH and LC-SPDP crosslinking B88 was used at pH 7.2, for SDAD crosslinking pH was 7.9. The total reaction volume for subsequent detection by immunoblotting was 100 µl with appropriate amount of crosslinker (SMPH or LC-SPDP: 1 mM; SDAD: 1.5 mM).

**Table 1.**
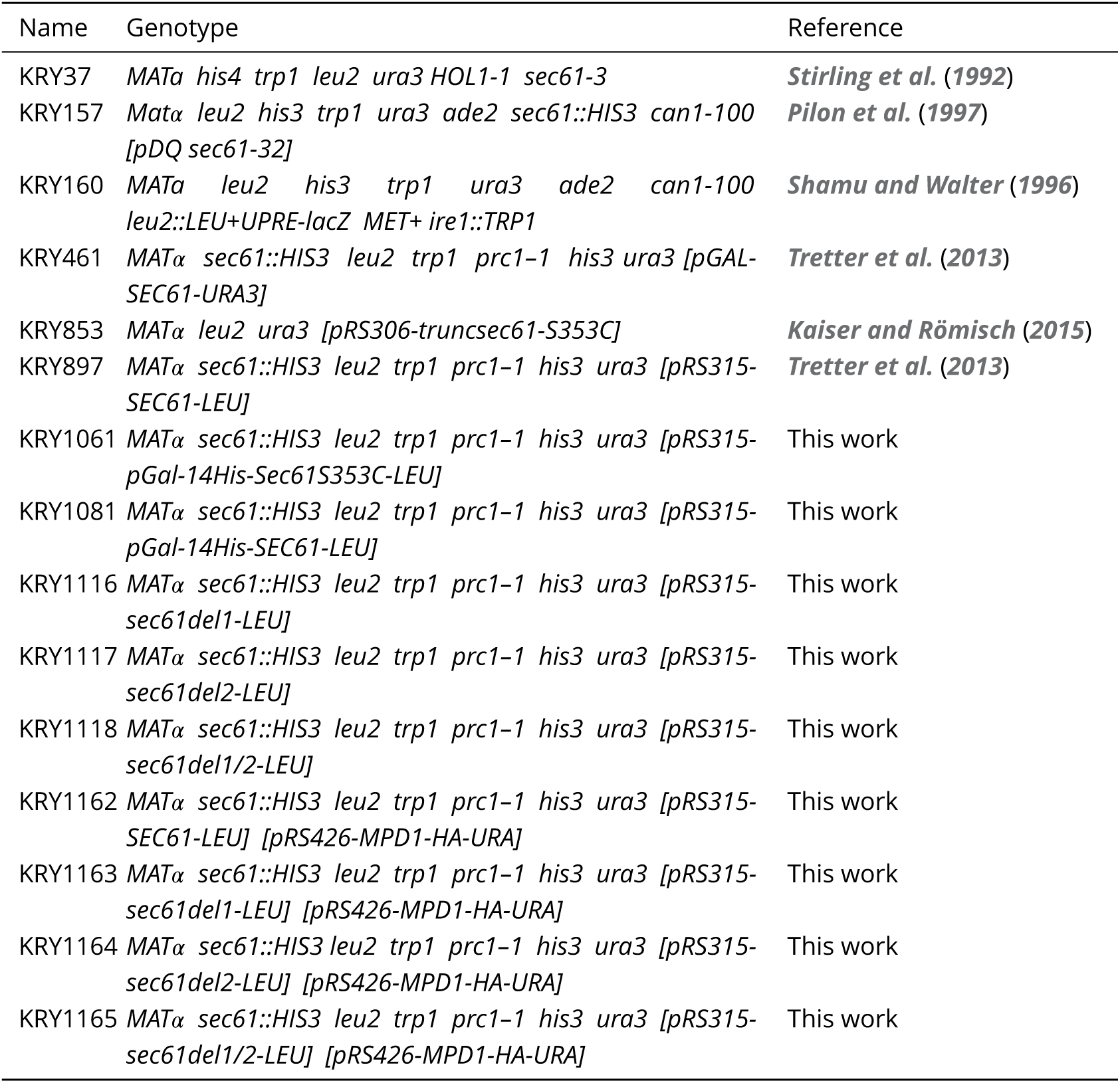
*S. cerevisiae* strains used in this study.

**Table 2.**
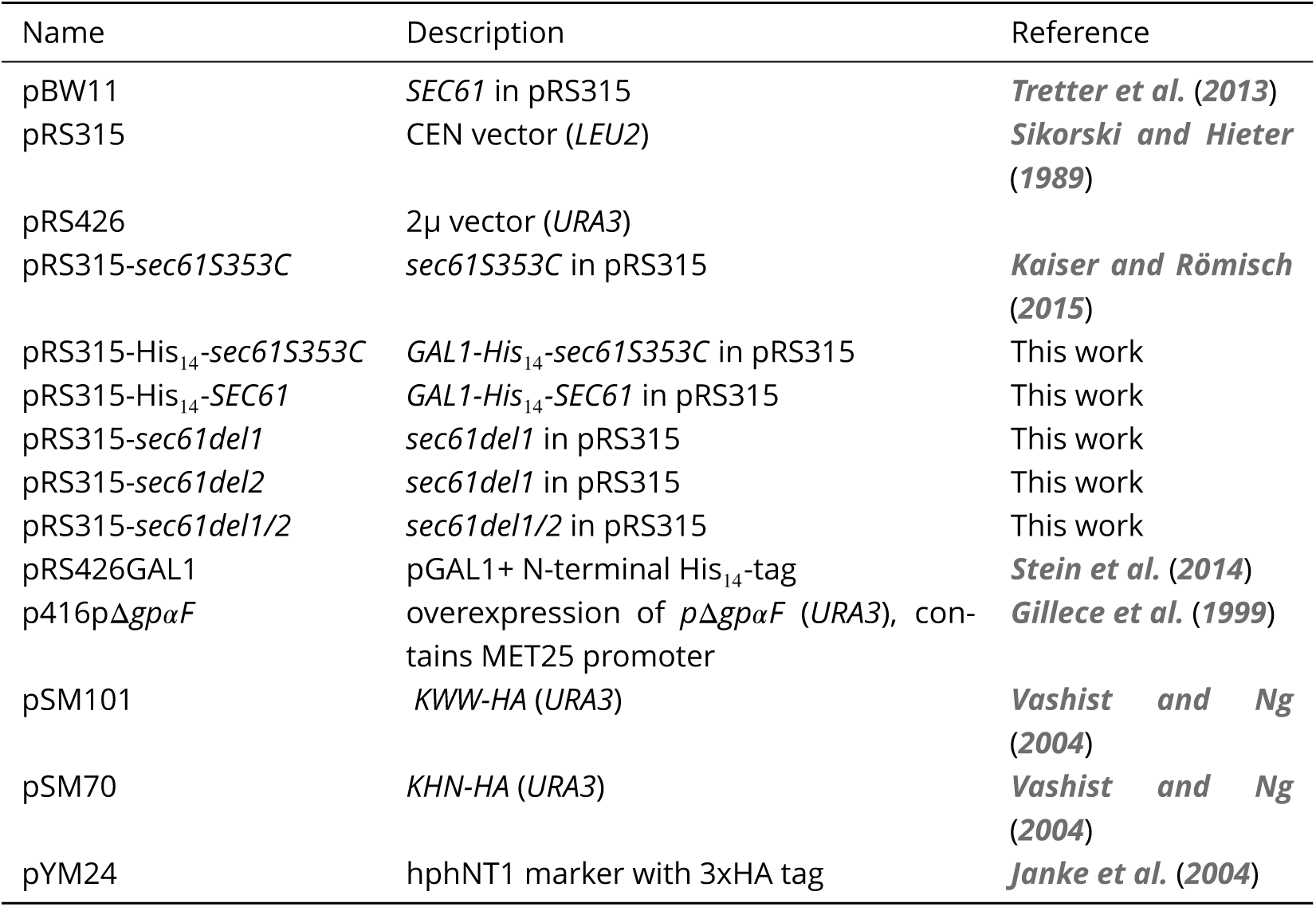
Plasmids used in this study.

**Table 3.**
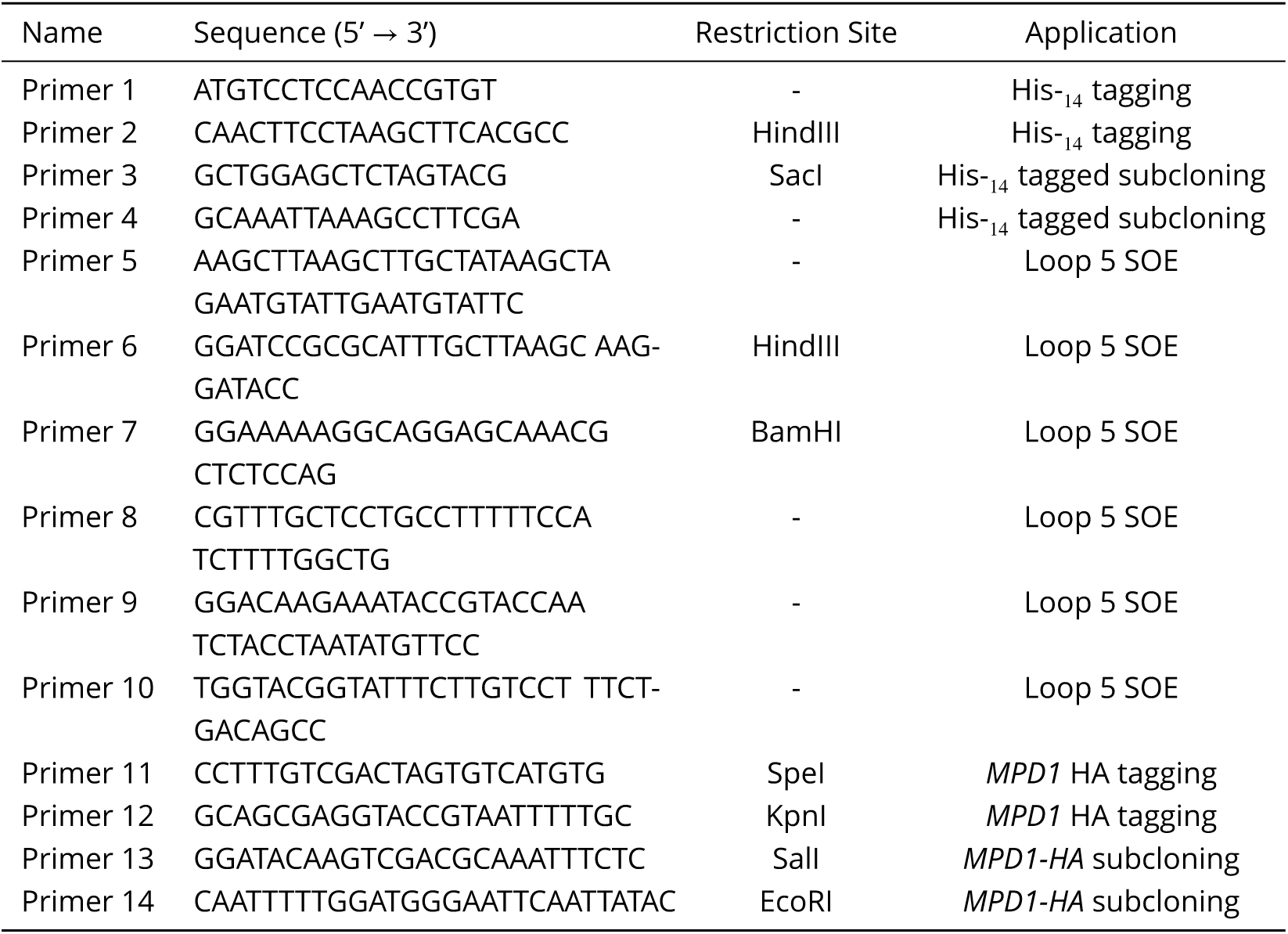
Primers used in this study.

**Table 4.**
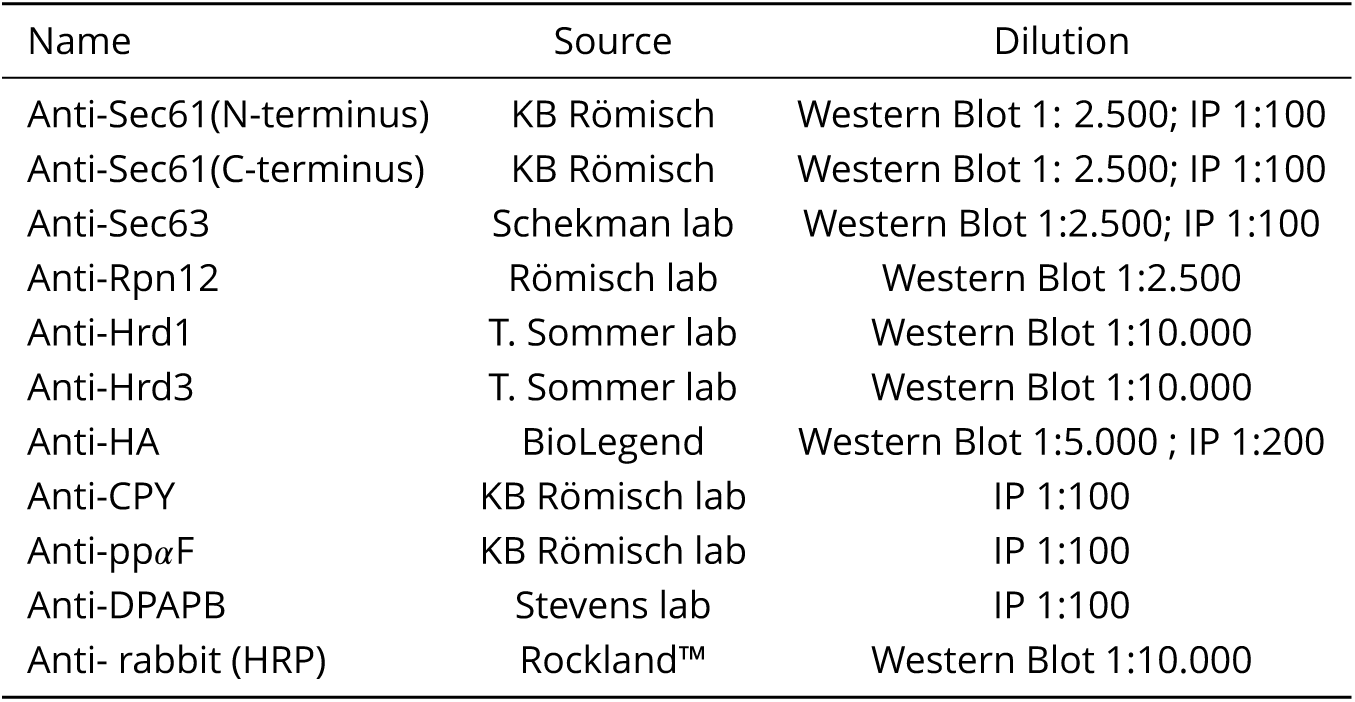
*S. cerevisiae* strains used in this study.

**Figure 1.**
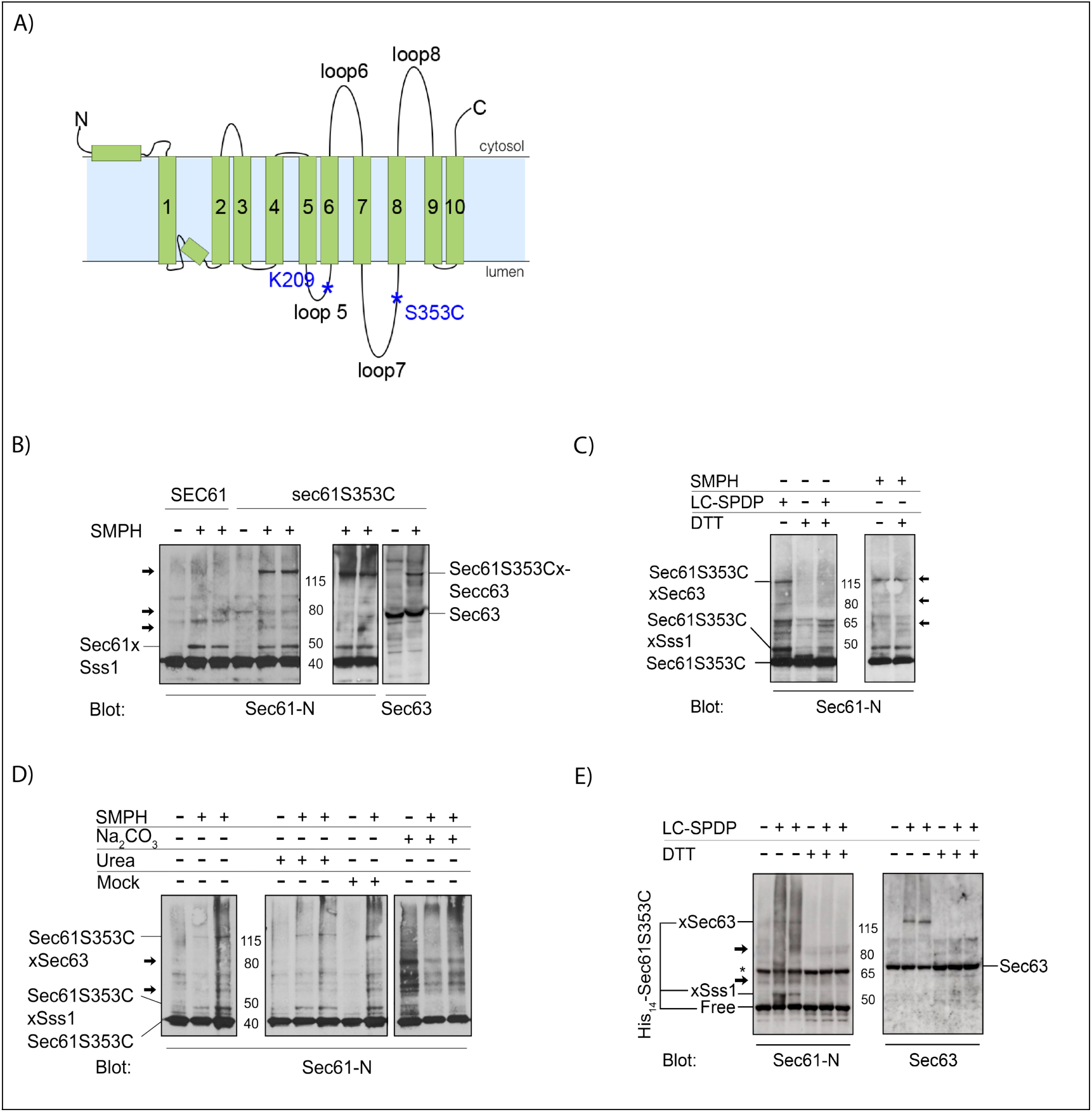
Optimization of crosslinking to Sec61S353C. A) Topological model of Sec61. **B)** Comparison of crosslinking patterns to Sec61 versus Sec61S353C with cysteine- and NH2-reactive SMPH. 17 eq microsomes per lane were crosslinked with 1mM SMPH on ice for 30 min and proteins resolved by SDS-PAGE. Sec61 was detected with an antibody against its N-terminus. Note that both Sec61 and Sec61S353C crosslink to Sss1. Additional crosslinked bands occurring in SecS353C samples are indicated by arrows in Sec61 panel. The largest product consists of Sec61S353C crosslinked to Sec63 (right panel). **C)** Sec61S353C crosslinking with SMPH (non-cleavable) or LC-SPDP (cleavable). Crosslinking was done as above and samples were resolved on SDS-PAGE without or with 200 mM DTT in the sample buffer as indicated. **D)** Crosslinking patterns to Sec61S353C after microsome extraction. Microsomes (17eq/lane) were extracted as indicated or mock-treated, crosslinked as above, and Sec61S353C and crosslinking products detected with an antibody against the Sec61 N-terminus. Note that crosslinks to Sec63 and Sss1 are sensitive to carbonate extraction. **E)** Crosslinking of His_14_-Sec61S353C microsomes with LC-SPDP. Crosslinking was done as above. Note that the N-terminal His_14_-tag did not affect crosslinking to Sec63 or Sss1 indicating no gross conformational alterations in the Sec complex. (*) indicates non-specific band that occurred independently of crosslinking in the Sec61 blot

**Figure 2.**
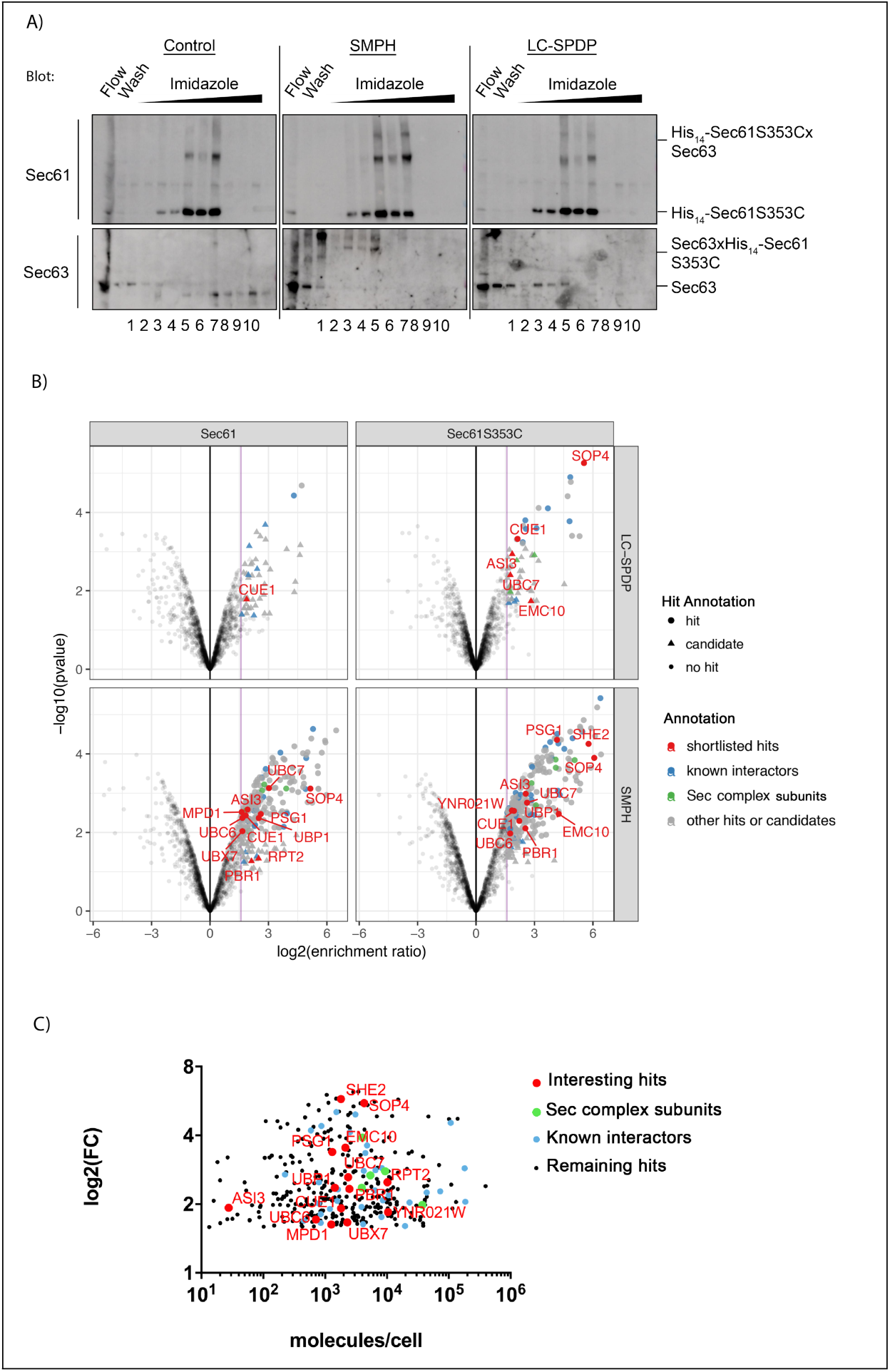
Purification and proteomics of Sec61-crosslinked proteins. A) 500 eq microsomes treated either with DMSO (control), SMPH (cleavable), or LC-SPDP (non-cleavable) were used. Samples were quenched and solubilized in IP Buffer. After denaturation (10 min, 65°C), proteins were diluted with cold Binding Buffer and applied to a HisTrap FF crude 1ml column. Fractionation was done using an imidazole gradient (100-500 mM). Sec61 and Sec63 were detected in each fraction after cleavage (LC-SPDP) and gel electrophoresis by immunoblotting with specific antibodies. **B)** Volcano plots based on the statistically determined protein enrichment in the crosslinked samples (His_14_-Sec61 and His_14_-Sec61S353C) when compared to the non-crosslinked samples. The horizontal axis represents log2 fold change (log2FC) reflecting level of enrichment. The vertical axis plots the -Log10(pValue) of enrichment, reflecting significance. Both hits and candidates have a fold change of at least 3. Hits have a false discovery rate (FDR) < 5 % and candidates an FDR < 20 %. Purple line is at fold-change of 3. Hits shown as colored dots and candidates as triangles. Elements of Sec61 complex in green; known interactors or translation machinery in blue; and shortlisted hits in red and points labeled on graph. Not significant hits below reference line and non-interesting hits above reference line in light grey. **C)** Graphical representation of the enrichment level (i.e logFC) of the Sec61 interactors as function of their respective cellular abundance as in Kulak et al. (2014). Known interactors blue, Sec61 complex subunits green, interesting interactors red and labeled on the graph. Note absence of correlation between cellular abundance and interaction with Sec61. **Figure 2 - Figure supplement 1** Table with mass spectrometry statistical analysis (hit and candidate determination).

**Figure 3.**
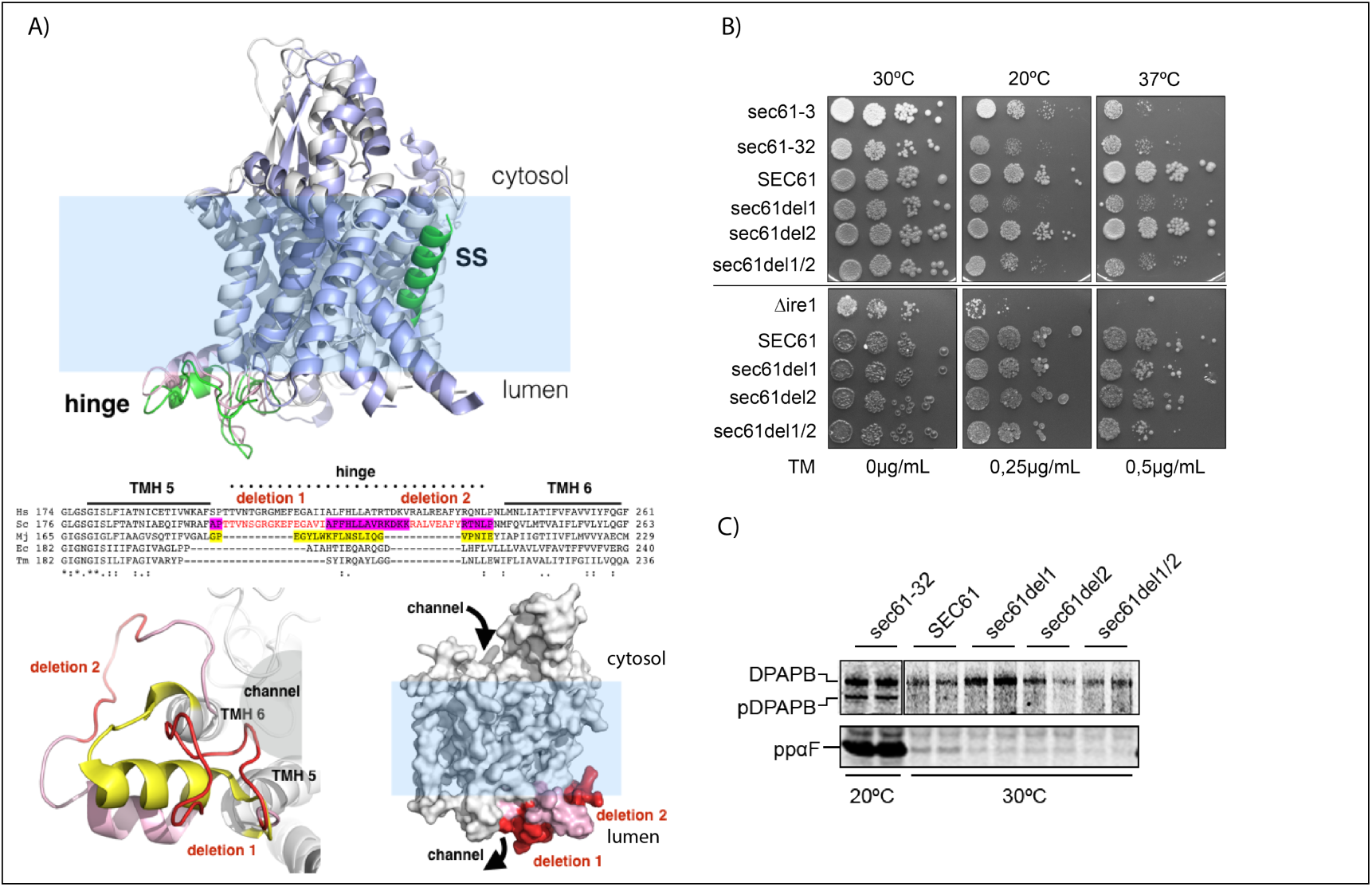
Design and characterization of *sec61* loop5 hinge mutants encompassing the binding site for Mpd1. A) Top: Structure of the Sec61 channel in closed (grey helices, pink hinge) versus open state (blue helices, green hinge, green signal sequence (SS) inserted in lateral gate) (Voorhees and Hegde, 2016) (PDB 3J7Q, PDB 3J7R). Note conformational changes in hinge (pink *vs* green) during channel opening. Middle: Alignment of loop5 hinge sequences of eukaryotes (*Homo sapiens*, Hs; *Saccharomyces cerevisiae*, Sc), prokaryotes (*Escherichia coli*, Ec; *Thermotoga maritima*, Tm) and archaea (*Methanococcus jannaschii*). Protein sequences were obtained from Uniprot. Regions coded by deletions in our *sec61* mutants are shown in red. The sequence forming the archaeal hinge region is highlighted in yellow, and the sequence corresponding to the vestigial (post-deletion) eukaryotic counterpart is highlighted in magenta. Bottom left: view of the hinge from the ER lumen (eukaryotic - PDB 3J7Q), showing the protein channel lined by TMHs 5, along with 6 and the intervening hinge (pink) with deletions 1 and 2 in red. The deletions result a shorter hinge akin to the archaeal structure shown in yellow (PDB 1RHZ) (also see middle). Bottom right: space filling model of Sec61 channel (PDB 3J7Q) in ER membrane indicating positions of deletions 1 and 2. Note that the region deleted in *sec61del1* is accessible for lumenal proteins in contrast to *sec61del2* which faces the membrane (lower right). **B)** Growth of *SEC61* and *sec61* hinge deletion mutants at different temperatures (30°C, 20°C, 37°C; top) or in the presence of tunicamycin (TM - 0 g/ml, 0.25 g/ml, 0.5 g/ml; all grown at 30°C, bottom). Cells (10^4^ to 10) were grown on YPD plates for 3 days. The following strains were used as controls: *sec61-3*, *sec61-32*, and Ll*ire1*. **C)** Analysis of ER import in *sec61* hinge mutants. Early log-phase cells were pulse labeled with [^35^S]-met/cys, lysed, and DPAPB (upper; cotranslational import) or prepro alpha-factor (pp*α*F) (lower; posttranslational import) immunoprecipitated. Starving and labeling were done at 30°C for all strains, except for *sec61-32*, which was incubated at 20°C. Labelling was done for 5 min for pp*α*F and 15 min for DPAPB. Proteins were detected by phosphorimaging. Figure 3 – Figuresupplement 1. *HAC1* mRNA Splicing Assay to evaluate UPR induction in *sec61* hinge mutants.

**Figure 4.**
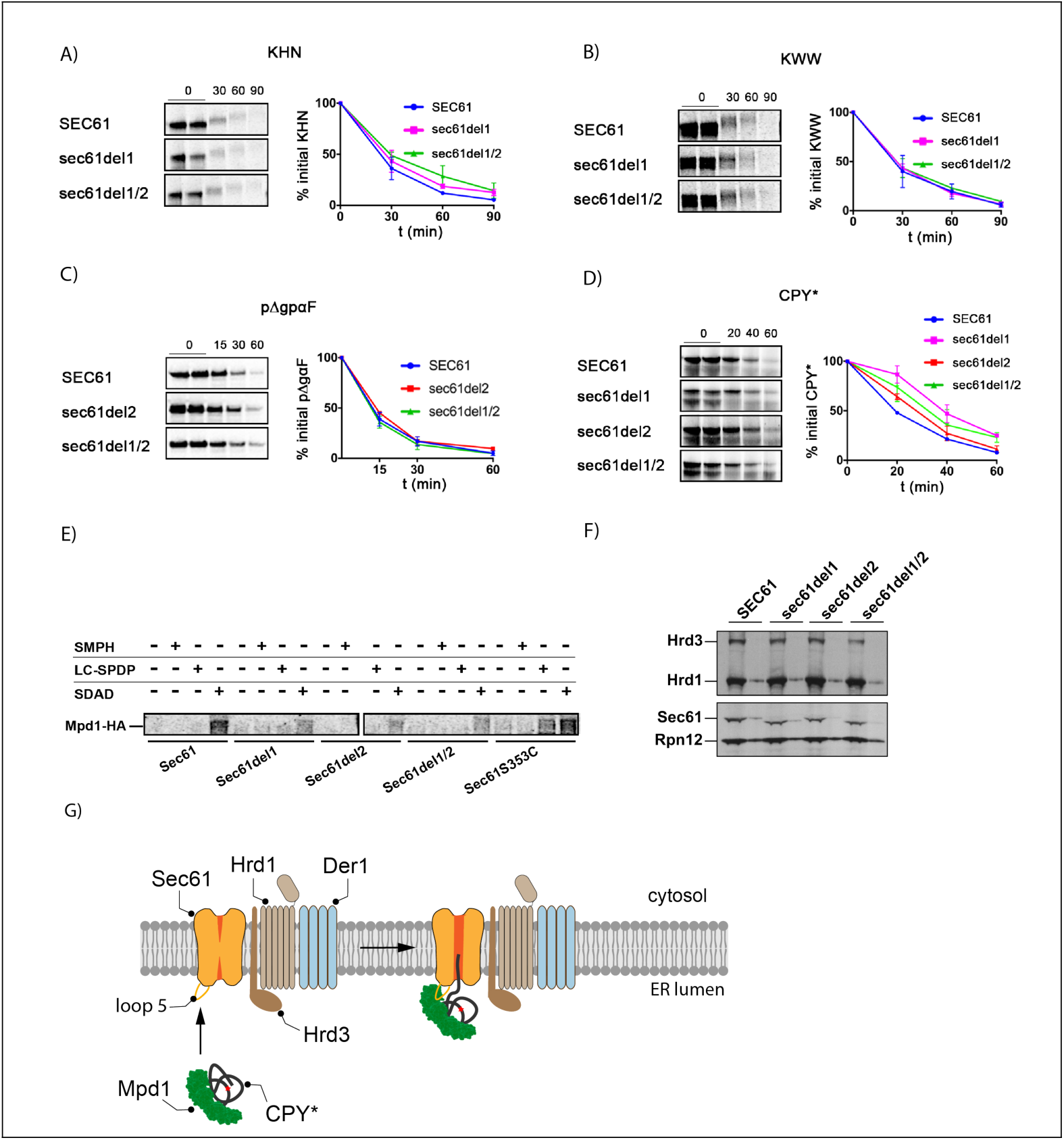
Mutation of the loop5 hinge in Sec61 specifically afects CPY* ERAD and interaction with Mpd1. A) - D) The *sec61* hinge mutants were screened for ERAD defects for: KWW; KHN, pΔgpαF, and CPY*. Wildtype and mutant strains were pulse-labeled with [^35^S]-met/cys for 5 (pΔgpαF and CPY*) or 15 min (KWW and KHN) followed by chase incubations for the indicated times. For each time point 1.5 OD_600_ of cells were lysed and proteins immunoprecipitated using specific antibodies (CPY*, pΔgpαF) or anti-HA. After SDS-PAGE, proteins were detected by phosphorimaging. Bands were quantified using ImageQuant (GE Healthcare) and averaged values plotted. For each experiment, at least three replicas were made. **E)** Interaction of Sec61 with Mpd1 was determined by crosslinking in [^35^S]-met/cys-labeled microsomes treated with SMPH (cysteine and NH_2_-reactive, non-cleavable), LC-SPDP (cysteine and NH_2_-reactive, cleavable) or SDAD (NH_2_-reactive and photoactivatable, cleavable) as indicated. For explanations of the crosslinker selection, see Material & Methods. Sec61 and crosslinked proteins were precipitated with anti-Sec61 N-terminal antibodies, followed by reduction of the crosslinker. Subsequently, Mpd1-HA was precipitated using HA-antibodies. After gel electrophoresis, proteins were detected by phosphorimaging. Equal amounts of cells were used for the preparation of each microsome batch. Protein levels of both Sec61 and Mpd1-HA were similar in all strains. Saturating amounts of antibodies were used for each precipitation. **F)** Steady state level of Sec61, Hrd1, and Hrd3 were determined by immunoblotting in wildtype and *sec61* hinge mutants. Two different amounts of cell lysates (1 and 1/3) of each sample were loaded side by side. Rpn12 was used as loading control. Specific antibodies for the different proteins were used for immublotting. **G)** Model for initiation of CPY* ERAD mediated by Mpd1 interaction with Sec61.

Control reactions were prepared with 5 µl of DMSO, but otherwise treated identically. For up-scaling, proportion of microsomes/total volume was maintained. After crosslinker addition, samples were incubated on ice for 30 min. Then, Quenching Buffer (1M Tris-HCl, pH 8; 100 mg/ml L-cys) was added (1/10 of total volume), and the sample incubated on ice for 15 min. Samples were then washed twice (always in the presence of quenching buffer) with appropriate pH B88, membranes sedimented at 16,000 x g for 10 min, and resuspended in appropriate form for subsequent use. For LC-SPDP cleavage, membranes were incubated for 15 min at room temperature in the presence of 100 mM of DTT. For SDAD crosslinking, after the washes the sample was exposed, on ice, to a 15 min UV (365 nm) irradiation with a 3UV Lamp (115V, 60Hz) (ThermoFisher) at a distance of 3,6 cm.

### Extraction of Luminal and Cytosolic Microsome-Associated Proteins

For extraction of cytosolic membrane-associated proteins, microsomes were resuspended in B88/Urea (20 mM Hepes-KOH, pH 6.8, 250 mM sorbitol, 150 mM KOAc, 5 mM Mg(OAc) _2_, 2,5 M urea), incubated for 20 min on ice, followed by sedimentation and washing of the membranes with B88, pH 6.8. For extraction of ER-luminal proteins, microsomes were resuspended in 100 mM sodium carbonate, pH 11.5, incubated on ice for 20 min, followed by sedimentation (20 min at 346,000xg, 4°C) of the membranes through a sucrose cushion (200 mM sucrose, 100 mM sodium carbonate, pH 11.5), and resuspension in B88, pH 6.8. For mock extractions, samples were treated in same way, but in absence of either urea or sodium carbonate.

### Immunoblotting

Protein gel electrophoresis was conducted using NuPAGE Novex pre-cast Bis-Tris gels (4–12,5% gels, 1.0 mm) and the XCell SureLock Mini-Cell (both Invitrogen). Proteins were transferred to nitrocellulose membranes (BioRad) and detected with specific antibodies at the appropriate dilutions, and an ECL reagent (Pierce) according to the supplier’s instructions. Signal was acquired either using an Amersham Imager 600 (GE Healthcare) or exposure to ECL films (Adavnsta).

### Purification of Sec61

ER membranes (500 eq) were treated as described in “Chemical Crosslinking”, either with DMSO (control), SMPH, or LC-SPDP in a total volume of 1.5 ml. After washing, membranes were resuspended in 150 µl of Quenching Buffer (1 M Tris-HCl, pH 8; 100 mg/ml L-cys) and diluted with 1 ml of IP Buffer (15 mM Tris-HCl, pH 7.5, 150 mM NaCl, 1 % Triton X-100, 0,1 % SDS) for solubilization (30 min at 4°C) followed by 10 min denaturation at 65°C. From this point on, all steps were done at 4°C. Sample was diluted with cold Binding Buffer (50 mM Tris-HCl, 300 mM KCl, 0,5 % Triton X- 100, 40 mM imidazole) to a final volume of 5 ml and applied to an HisTrap FF crude (1 ml) column integrated into a BioLogic automated purification system (Biorad). After sample loading (0.5 ml/min for 10 ml), the column was washed with Binding Buffer (10 ml; 1 ml/min) and sample eluted along a step gradient of imidazole (100-500 mM, 15 ml per step, 1ml/min. Steps: 100; 200; 400; 500). Fractions (7,5 ml) were collected along the gradient with an automatic fraction collector. DTT (100 mM) was added to each fraction. Each differently treated sample was applied to an independent column. Between purifications, the system was washed with 10 ml H_2_O, 10 ml ethanol 20 %, 10 ml H_2_O, 20 ml Binding Buffer. Fractions where Sec61 was eluted (fraction 3-10 - 50 ml total) were pooled, proteins precipitated with 10 % TCA on ice for 2h and washed with ice-cold acetone. Each pellet was resuspended in 2 x Laemmli Buffer, and resolved for 5 cm on 4-12,5% NuPAGE gel. The gel was then stained by Coomassie Colloidal Staining (0.08% Coomassie Brilliant Blue G250 (CBB G250), 10 % citric acid, 8% ammonium sulfate, 20 % methanol) overnight and destained with water as described in the EMBL online Proteomics Core Facility Protocols. The gels where then sealed in individual plastic bags with a few milliliters of water and shipped to the Mass Spectrometry Facility.

### Mass Spectrometry

#### Sample preparation

The whole lane of each samples was cut out into small cubes and subjected to in-gel digestion with trypsin (Savitski et al., 2014). After overnight digestion, peptides were extracted from the gel pieces by sonication for 15 minutes, tubes were centrifuged, the supernatant removed and placed in a clean tube. Followed by a second extraction round with a solution of 50:50 water: acetonitrile, 1% formic acid (2 x the volume of the gel pieces) and the samples were sonicated for 15 minutes, centrifuged and the supernatant pooled with the first extract. The pooled supernatants were then subjected to speed vacuum centrifugation. Samples were reconstituted in 96:4 water: acetonitrile, 0.1% formic acid and further processed using an OASIS^®^ HLB µElution Plate (Waters) according the manufacturer’s instructions.

#### LC-MS/MS

Peptides were separated using the nanoAcquity Ultra Performance Liquid Chromatography (UPLC) system (Waters) using a trapping (nanoAcquity Symmetry C18, 5 µm, 180 µm x 20 mm) as well as an analytical column (nanoAcquity BEH C18, 1.7 µm, 75 µm x 200 mm). The outlet of the analytical column was coupled to a Linear Trap Quadrupole (LTQ) Orbitrap Velos Pro (Thermo Fisher Scientific) using the Proxeon nanospray source. Solvent A consisted of water, 0.1% formic acid and solvent B consisted of acetonitrile, 0.1% formic acid. Sample was loaded with a constant flow of solvent A at 5 µl/min onto the trapping column. Peptides were eluted over the analytical column with a constant flow of 0.3 µl/min During elution the percentage of solvent B increased linearly from 3% to 7% in 10 min., then increased to 25% in 110 min and to 40% for the final 10 min a cleaning step was applied for 5 min with 85% B followed by 3% B 20 min. The peptides were introduced into the mass spectrometer via a Pico-Tip Emitter 360 µm OD x 20 µm ID; 10 µm tip (New Objective), a spray voltage of 2.2 kV was applied. Capillary temperature was 300 °C. Full scan MS spectra were acquired with a resolution of 30000. The filling time was set at a maximum of 500 ms with a maximum ion target of 1.0 x 10^6^. The fifteen most intense ions from the full scan MS (MS1) were sequentially selected for sequencing in the LTQ. Normalized collision energy of 40% was used, and the fragmentation was performed after accumulation of 3.0 x10^4^ ions or after a maximum filling time of 100 ms for each precursor ion (whichever occurred first). Only multiply charged (2^+^, 3^+^, 4^+^) precursor ions were selected for MS/MS. The dynamic exclusion list was restricted to 500 entries with maximum retention period of 30 s and a relative mass window of 10 ppm. In order to improve the mass accuracy, a lock mass correction using the ion (m/z 445.12003) was applied.

#### Data analysis

The raw mass spectrometry data was processed with MaxQuant (v1.5.2.8) (Cox and Mann, 2008) and searched against an Uniprot Saccharomyces cerevisiae proteome database. The search parameters were as follows: Carbamidomethyl (C) (fixed), Acetyl (N-term) and Oxidation (M) (variable) were used as modifications. For the full scan MS spectra (MS1) the mass error tolerance was set to 20 ppm, and for the MS/MS spectra (MS2) to 0.5 Da. Trypsin was selected as protease with a maximum of two missed cleavages. For protein identification a minimum of one unique peptide with a peptide length of at least seven amino acids and a false discovery rate below 0.01 were required on the peptide and protein level. The match between runs function was enabled, a time window of one minute was set. Label free quantification was selected using iBAQ (calculated as the sum of the intensities of the identified peptides and divided by the number of observable peptides of a protein) (Schwanhäusser et al., 2011) with the log fit function enabled.

We also used the xQuest/xProphet pipeline (Leitner et al., 2014) to identify crosslinked peptides in our samples. For this, we used the basic protocol and conditions used in Leitner et al. (2014), correcting the meaningful parameters to fit our crosslinker (e.g monoisotopic shift, only light chain, reactive groups, etc.). Databases of no more than 30 proteins were fed into the pipeline.

#### Statistical Analysis

The raw output data of MaxQuant (proteinGroups.txt file) was processed using the R programming language (ISBN 3-900051-07-0). As a quality filter we allowed only proteins that were quantified with at least 2 unique peptides. Potential batch-effects were removed from the log2 of the iBAQ values using the limma package (Ritchie et al., 2015). Furthermore, batchcleaned data were normalized with the vsn package (variance stabilization) (Huber et al., 2002). Missing values were imputed using the MSNbase package (Gatto and Lilley, 2012). For conditions with at least 2 out of 3 identifications, the “knn” method was used. For less identifications, the “MinDet” method was applied. Finally, limma was used again to identify differentially expressed proteins. A protein was called a hit with a false discovery rate (fdr) smaller 5 % and a fold change of at least 3 and a candidate with a fdr smaller 20 % and a fold change of at least 3.

### Mutant Construction

#### 14His-Tagged constructs

For His_14_-tagging of *SEC61* and *sec61S353C*, both genes were amplified from pBW11 and pRS315- *sec61S353C*, respectively, using Primer 1 and Primer 2. The resulting PCR products were cloned into pRS426pGAL1 (Stein et al., 2014) using the SfoI and HindIII restriction sites. Correct cloning was confirmed by sequencing. The pGal-His_14_-*SEC61*-CYC and pGal-His_14_-*Sec61S353C*-CYC cassettes were then amplified using Primer 3 and Primer 4. The resulting PCR products were cloned into pRS315 (CEN, LEU2). Transformants in the JDY638 (pGAL-SEC61-URA3) *S. cerevisiae* background were first selected on SC -URA medium containing 2% (w/v) galactose and 0.2% (w/v) glucose lacking leucine. The pGAL-SEC61 plasmid was selected against on SC 5-FOA plates containing 2% (w/v) galactose and 0.2% (w/v) glucose without leucine. Constructs were confirmed by sequencing.

#### *SEC61* Loop 5 deletion mutants

Mutants *sec61del1*, *sec61del2*, and *sec61del1/2* were generated by PCR-driven overlap extension (SOE PCR) (Aiyar et al., 1996; Horton et al., 1989) followed by transformation into KRY461 of the respective constructs. For the initial SOE-PCR reactions, *SEC61* was amplified from pBW11 (Table 2). Deletion 1 and deletion 2 were made separately. Deletion 1/2 was made using deletion 1 construct as template and same primers as used for the generation of deletion 2. For SOE-PCR, the regions upstream and the downstream of the deletion sites were amplified using a mutagenic primer and a gene flanking primer (Table 3). Each mutagenic primer immediately flanks the deletion site and both upstream and downstream deletion-flanking primer have a stretch of complementarity with each other. For the extension of the final PCR product, the gene-flanking primer-pair was used and both upstream and downstream fragments were used as template (working as a single-template unit). The resulting PCR products were cloned into pRS315 (CEN, *LEU2*) (Sikorski and Hieter, 1989). Transformants into JDY638 (*pGAL-SEC61-URA3*) were first selected on SC -URA medium containing 2% (w/v) galactose and 0.2% (w/v) glucose without leucine. The *pGal-SEC61* plasmid shuffle was done on SC 5’-FOA plates lacking leucine. All constructs were confirmed by sequencing.

#### *MPD1* HA-Tagging

Tagging of genomic *MPD1* was done as described in Janke et al. (2004). Briefly, the HA cassette was amplified from pYM24 (supplied by Michael Knop) using Primer11 and Primer12. The plasmid contains the HA-cassette as well as the hphNT1 for selection. Targeting was done by homology of the designed primers with the appropriate regions of the gene of interest. This PCR product was then used to transform KRY461, and transformants were selected on YPD plates containing hygromycin (300µg/ml). *MPD1-HA* was amplified from the genomic DNA using Primer13 and Primer14 and cloned into pRS426 (2µM, *URA3*). This plasmid was then used to transform the hinge mutant strains.

### Cell Labelling and Immunoprecipitation

Aliquots of 1.5 OD_600_ early log phase cells were incubated in synthetic media lacking methionine, cysteine, and ammonium sulfate for 15 or 30 min (depending on the protein to be labelled) at the appropriate temperature and shaking at 220 rpm. Cells were labeled with [^35^ S]-met/cys (Express Labeling, PerkinElmer) (1.5 MBq per sample) mix for 5 min (CPY*, pΔgp*α*F) or 15 min (DPAPB, KWW, KHN). For pulse experiments, after labeling cells were immediately killed with Tris-Azide Buffer (20 mM Tris-HCl, pH 7.5, 20 mM sodium azide). For pulse-chase experiments, zero time points were treated as above, and to remaining samples Chase Mix (0.03% cys, 0.04% met, 10 mM ammonium sulfate) was added, and samples were incubated with shaking at the appropriate temperature for the indicated times. At each time point, Tris-Azide Buffer was added. Cells were harvested and incubated in 100 mM Tris-HCl, pH 9.4, for 10 min at room temperature. Subsequently, samples were lysed with glass beads in Lysis Buffer (20 mM Tris-HCl, pH 7.5, 2% (w/v) SDS, 1 mM DTT, 1 mM PMSF) and denatured for 5 min at 95°C (soluble proteins) or 10 min at 65°C (transmembrane proteins). Afterwards, glass beads were washed 3 times and the combined washes used for immunoprecipitation after preclearing with 60 µl 20% Protein A-Sepharose beads (GE Healthcare) in IP-buffer (15 mM Tris-HCl, pH 7.5, 150mM NaCl, 1% Triton X-100, 0,1% SDS) (Pilon et al., 1997). Precipitations were done with 60 µl 20% Protein A-Sepharose beads (GE Healthcare) and appropriate amount of antibody, either at room temperature for 2h or at 4°C for 4h or over night. Protein A-Sepharose beads were washed as in Baker et al. (1988), proteins eluted with 2x Laemmli Buffer and denatured at 95°C for 5 min (soluble) or 65°C for 10 min (transmembrane). Proteins were resolved on 4-12,5% NuPAGE gels. Dried gels were exposed to Phosphorimager plates, and the signal acquired with a Typhoon PhosphoImager (GE Healthcare).

### Detection of Sec61 Interactors in Radiolabeled Membranes

Crude radiolabeled ER vesicles (10 µl) were crosslinked as described in “Chemical Crosslinking” and submitted to two consecutive immunoprecipitations. Hinge mutants are derived from a *SEC61* background. Microsomes from the *sec61S353C* strain were included, because Sec61-Mpd1 interaction was first detected in this strain. Crosslinker selection: The Sec61-Mpd1 crosslinked peptide was first identified by SMPH crosslinking to Sec61S353C. SMPH and LC-SPDP have one cysteine- and one NH_2_-reactive group. Only LC-SPDP is cleavable, so in the double immunoprecipitation experiment, SMPH is teh negative control for LC-SPDP, because there should be no release of Mpd1 from Sec61 after the first precipitation. SDAD is also cleavable, but with one NH_2_ -reactive and one photoactivatable reactive group. It was used to effciently crosslinke Mpd1 to Sec61 regardless of the cysteine in loop 7. For the first precipitation, the membranes were solubilized in Lysis Buffer (20 mM Tris, pH 7.5, 2% SDS, 1 mM PMSF) and denatured at 65°C for 10 min. Proteins were then diluted in Washing Buffer (15 mM Tris-HCl, pH 7.5, 150 mM NaCl, 1% Triton X-100, 2 mM NaN3, 1 mM PMSF). After pre-clearing (as previously), 60 µl of 20% Protein A-Sepharose beads (GE Healthcare) and appropriate amount of Sec61 antibody was added. Samples were then incubated with rotation overnight at 4°C, and Protein A Sepharose pellets washed as above. For elution we used 20 µl of 20 mM Tris-HCl, pH 7.5, 5% SDS, 50 mM DTT for 15 min room temperature and denaturation for 10 min 65°C. Eluted proteins were then diluted in Washing Buffer and the Mpd1-HA precipitated using anti-HA polyclonal antibody (BioLegend). Precipitation was done for 2h at room temperature followed by elution done 2 x Laemmli Buffer, 200 mM DTT. Proteins were denatured again as before, resolved on 4-12,5% NuPAGE gels exposed to Phosphorimager plates, and the signal acquired with a Typhoon PhosphoImager (GE Healthcare).

## Acknowledgments

We thank Rainer Pepperkok (EMBL Heidelberg) for access to the EMBL Mass Spectrometry facility; Alexander Leitner (ETH Zürich) for help with xQuest/xProphet and data interpretation; Alexander Stein and Tom Rapoport (Howard Hughes Medical School), Randy Schekman (UC Berkeley), Randy Hampton (UC San Diego), Thomas Sommer & Ernst Jarosch (MDC Berlin), Davis Ng (TLL Singapore), Michael Knop (ZMBH Heidelberg), Pedro Carvalho (Oxford University) for antibodies and plasmids; Maya Schuldiner (Weizmann Institute) for advice on identifying interesting Sec61 interactors and unpublished information about several of these; Timo Scheller (Saarland University IT) for helping with xQuest/xProphet pipeline setup; Carmen Clemens for technical assistance.

## Contributions

Contributions for this paper were as follow:

1. Fábio Pereira performed and conceived experiments, analyzed results, and wrote the manuscript.
2. Mandy Rettel and Mikhail M. Savitski performed mass spectrometry experiments.
3. Frank Stein performed the statistical analysis of all mass spectrometry data.
4. Ian Collinson designed deletion mutants in loop5 of Sec61.
5. Karin Römisch conceived experiments, analyzed results, wrote the manuscript.

## Supplemental Figures

**Figure 3 – Figure supplement 1.**
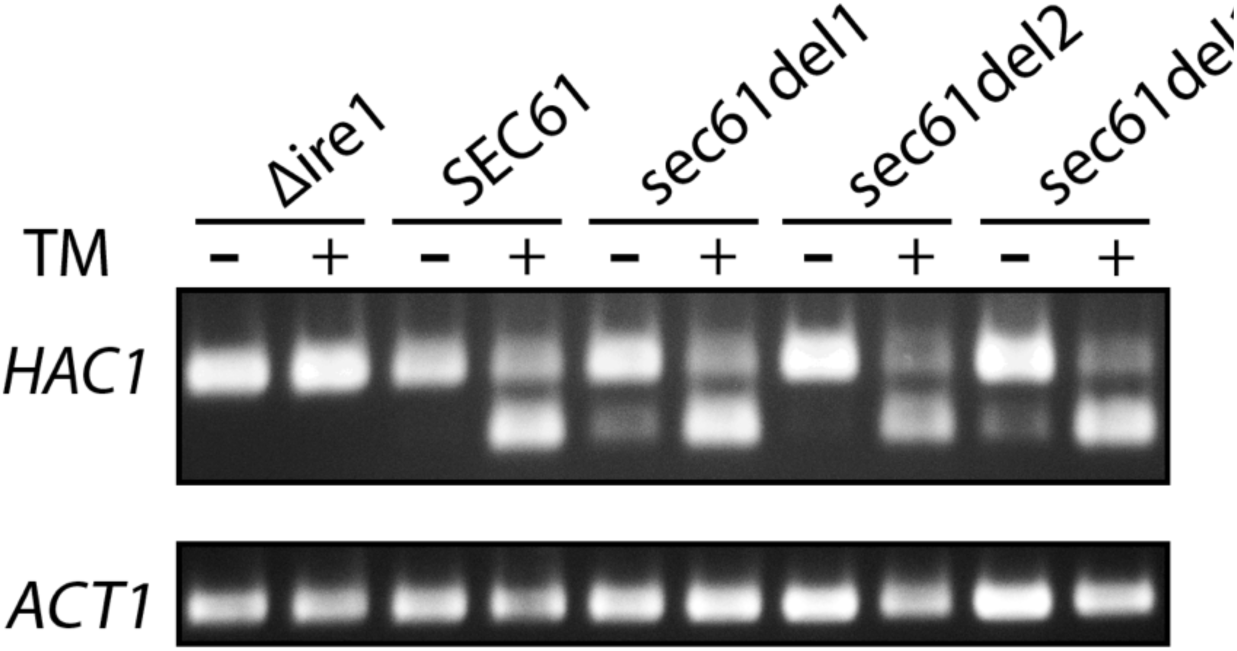
– Figure supplement 1 –. *HAC*1 mRNA Splicing Assay to evaluate UPR induction. Wildtype and Sec61 hinge mutants were either treated with tunicamycin (2 µg/ml) (TM) or DMSO (control), followed by total RNA isolation, and cDNA production from isolated RNA. A quantitative PCR was done from equal amounts of cDNA. Agarose gel showing the resultant PCR products. Upper slice shows *HAC1* PCR product. Upper bands (720 bp) represent the unspliced (uninduced) *HAC1* mRNA, while lower bands (470 bp) represent the spliced (induced) *HAC1* mRNA. Bottom slice show the actin PCR product. The *?ire1* mutant was used as negative control.

## Supplemental Methods

### RNA isolation and HAC splicing PCR

For the isolation of RNA all solutions were RNAse free. Strains to be evaluated were grown to an OD_600_=1, and two 10 ml replicas per strain were made. To one replica tunicamycin (2 µg/ml of) was added, to the other DMSO (same volume as tunicamycin), and cells were grown for 3h more. Cells were then harvest at 4,500 x g for 5 min (4°C), resuspended in 1 ml ice-cold DEPC- water, and transferred to an RNase-free tube. After sendimentation (13,000 x g, 10 sec, 4°C) pellet was resuspended in 400 µl TES Solution (10 mM Tris-HCl, pH 7.7, 10 mM EDTA, 0.5% (w/v) SDS), 400 µl of Roti-Aqua-Phenol^®^ (Carl Roth) were added, and after vortexing (10 sec), samples were incubated for 1 h at 65°C with occasional vortexing. Samples were then placed on ice for 5 min and centrifuged at 13,000 x g for 5 min (4°C). Aqueous phase was transferred to a clean tube and 400 µl of Roti-Aqua-Phenol^®^ were added. Samples were vortexed for 20 sec and incubated for 5 min on ice. Samples were then centrifuged as before, aqueous phase transferred again to a clean tube, and 400 µl of chloroform were added. Samples were vortexed again (20sec) and sendimented (13,000 x g, 5 min, 4°C). Aqueous phase was once more transferred to a clean tube, and 40 µl of 3M NaAc, followed by 1 ml of ice cold 100% ethanol, were added. After repeating the vortexing and sedimentation steps, pellets were washed with 1.5 ml of 70% ethanol and sedimented as before. Finally, samples were resuspended in 50 µL of DEPC-water and RNA concentration was determined using a NanoDrop spectrophotometer (ThermoFisher).

To generate cDNA from each RNA samples, the RNA samples were diluted to a concentration of 0.1 µg/ml and reverse-transcription reactions were made as follows using MaximaRT^®^ (ThermoFisher):

| Component | Volume (μl) | Final concentration |
| --- | --- | --- |
| RNA | 1 | 0.1 μg |
| Oligo(dT18)-primer (100 mM) | 1 | 100 pmol |
| dNTP mix (10 mM) | 1 | 0.5 mM |
| RNase-free dH<sub>2</sub>O | To 14.5 | to 14.5 μl |
| 5X RT buffer | 4 | 1x |
| RNasin (40 U/μl) | 0.5 | 20 U |
| Maxima® RT | 1 | 200 U |

Samples were then incubated for 30min at 50°C followed by an inactivation at 85°C for 5 min. We then used 1 µl of each cDNA for PCR, using both the *HAC1*- (5’- CTGGCTGACCACGAAGAC and 5’- TTGTCTTCATGAAGTGATGGC-3’) and the *ACT1*- (5’- ATTCTGAGGTTGCTGCTTT-3’ and 5’- GTGGTGAACGATAGATGG-3’) specific primers.

Amplification reactions were done using KAPAHiFi™ Hot Start DNA (PEQLAB) and the program used was the following:

| Cycles | Step | Temperature | Duration |
| --- | --- | --- | --- |
| 1 | Initial denaturation | 95 | 5 sec |
| 35 | Denaturation | 98 | 20 sec |
|  | Annealing | 54 | 15 sec |
|  | Extension | 72 | 30 sec/kb |
| 1 | Final Extension | 72 | 5 min |
|  | Store | 4 | ∞ |

After PCR 10 µl of each reaction was resolved in an 1% agarose gel at 100V for 1h. Signal was acquired with the E-BOX VX2 gel documentation system (PEQLAB).

## References

Aiyar A, Xiang Y, Leis J. Site-directed mutagenesis using overlap extension PCR. Methods Mol Biol. 1996; 57:177–91. doi: 10.1385/0-89603-332-5:177.

Aviram N, Ast T, Costa EA, Arakel EC, Chuartzman SG, Jan CH, Haßdenteufel S, Dudek J, Jung M, Schorr S, Zimmermann R, Schwappach B, Weissman JS, Schuldiner M. The SND proteins constitute an alternative targeting route to the endoplasmic reticulum. Nature. 2016 11; 540(7631):134–138. doi: 10.1038/nature20169.

Baker RT, Tobias JW, Varshavsky A. Ubiquitin-specific proteases of Saccharomyces cerevisiae. Cloning of UBP2 and UBP3, and functional analysis of the UBP gene family. J Biol Chem. 1992 Nov; 267(32):23364–75.

Brodsky JL, Goeckeler J, Schekman R. BiP and Sec63p are required for both co- and posttranslational protein translocation into the yeast endoplasmic reticulum. Proc Natl Acad Sci U S A. 1995 Oct; 92(21):9643–6.

Carvalho P, Goder V, Rapoport TA. Distinct ubiquitin-ligase complexes define convergent pathways for the degradation of ER proteins. Cell. 2006 Jul; 126(2):361–73. doi: 10.1016/j.cell.2006.05.043.

Christianson JC, Olzmann JA, Shaler TA, Sowa ME, Bennett EJ, Richter CM, Tyler RE, Greenblatt EJ, Harper JW, Kopito RR. Defining human ERAD networks through an integrative mapping strategy. Nat Cell Biol. 2011 Nov; 14(1):93–105. doi: 10.1038/ncb2383.

Cox J, Mann M. MaxQuant enables high peptide identification rates, individualized p.p.b.-range mass accuracies and proteome-wide protein quantification. Nat Biotechnol. 2008 Dec; 26(12):1367–72. doi: 10.1038/nbt.1511.

Esnault Y, Feldheim D, Blondel MO, Schekman R, Képès F. SSS1 encodes a stabilizing component of the Sec61 subcomplex of the yeast protein translocation apparatus. J Biol Chem. 1994 Nov; 269(44):27478–85.

Foresti O, Rodriguez-Vaello V, Funaya C, Carvalho P. Quality control of inner nuclear membrane proteins by the Asi complex. Science. 2014 Nov; 346(6210):751–5. doi: 10.1126/science.1255638.

Gatto L, Lilley KS. MSnbase-an R/Bioconductor package for isobaric tagged mass spectrometry data visualization, processing and quantitation. Bioinformatics. 2012 Jan; 28(2):288–9. doi: 10.1093/bioinformatics/btr645.

Ghaemmaghami S, Huh WK, Bower K, Howson RW, Belle A, Dephoure N, O’Shea EK, Weissman JS. Global analysis of protein expression in yeast. Nature. 2003 Oct; 425(6959):737–41. doi: 10.1038/nature02046.

Gillece P, Luz JM, Lennarz WJ, de La Cruz FJ, Römisch K. Export of a cysteine-free misfolded secretory protein from the endoplasmic reticulum for degradation requires interaction with protein disulfide isomerase. J Cell Biol. 1999 Dec; 147(7):1443–56.

Grubb S, Guo L, Fisher EA, Brodsky JL. Protein disulfide isomerases contribute differentially to the endoplasmic reticulum-associated degradation of apolipoprotein B and other substrates. Mol Biol Cell. 2012 Feb; 23(4):520–32. doi: 10.1091/mbc.E11-08-0704.

Horton RM, Hunt HD, Ho SN, Pullen JK, Pease LR. Engineering hybrid genes without the use of restriction enzymes: gene splicing by overlap extension. Gene. 1989 Apr; 77(1):61–8.

Huber W, von Heydebreck A, Sültmann H, Poustka A, Vingron M. Variance stabilization applied to microarray data calibration and to the quantification of differential expression. Bioinformatics. 2002; 18 Suppl 1:S96–104.

Jadhav B, McKenna M, Johnson N, High S, Sinning I, Pool MR. Mammalian SRP receptor switches the Sec61 translocase from Sec62 to SRP-dependent translocation. Nat Commun. 2015 Dec; 6:10133. doi: 10.1038/ncomms10133.

Janke C, Magiera MM, Rathfelder N, Taxis C, Reber S, Maekawa H, Moreno-Borchart A, Doenges G, Schwob E, Schiebel E, Knop M. A versatile toolbox for PCR-based tagging of yeast genes: new fluorescent proteins, more markers and promoter substitution cassettes. Yeast. 2004 Aug; 21(11):947–62. doi: 10.1002/yea.1142.

Johnson AE, van Waes MA. The translocon: a dynamic gateway at the ER membrane. Annu Rev Cell Dev Biol. 1999; 15:799–842. doi:10.1146/annurev.cellbio.15.1.799.

Kaiser ML, Römisch K. Proteasome 19S RP binding to the Sec61 channel plays a key role in ERAD. PloS One. 2015; 10(2):e0117260. doi:10.1371/journal.pone.0117260.

Kalies KU, Görlich D, Rapoport TA. Binding of ribosomes to the rough endoplasmic reticulum mediated by the Sec61p-complex. J Cell Biol. 1994 Aug; 126(4):925–34.

Kalies KU, Rapoport TA, Hartmann E. The beta subunit of the Sec61 complex facilitates cotranslational protein transport and interacts with the signal peptidase during translocation. J Cell Biol. 1998 May; 141(4):887–94.

Kulak NA, Pichler G, Paron I, Nagaraj N, Mann M. Minimal, encapsulated proteomic-sample processing applied to copy-number estimation in eukaryotic cells. Nat Methods. 2014 Mar; 11(3):319–24. doi:10.1038/nmeth.2834.

Leitner A, Walzthoeni T, Aebersold R. Lysine-specific chemical cross-linking of protein complexes and identifica- tion of cross-linking sites using LC-MS/MS and the xQuest/xProphet software pipeline. Nat Protoc. 2014 Jan; 9(1):120–37. doi:10.1038/nprot.2013.168.

Mehnert M, Sommer T, Jarosch E. Der1 promotes movement of misfolded proteins through the endoplasmic reticulum membrane. Nat Cell Biol. 2014 Jan; 16(1):77–86. doi:10.1038/ncb2882.

Neal S, Jaeger PA, Duttke SH, Benner C, K Glass C, Ideker T, Hampton RY. The Dfm1 Derlin Is Required for ERAD Retrotranslocation of Integral Membrane Proteins. Mol Cell. 2018 Jan; 69(2):306–320.e4. doi:10.1016/j.molcel.2017.12.012.

Ng W, Sergeyenko T, Zeng N, Brown JD, Römisch K. Characterization of the proteasome interaction with the Sec61 channel in the endoplasmic reticulum. J Cell Sci. 2007 Feb; 120(Pt 4):682–91. doi:10.1242/jcs.03351.

Pilla E, Schneider K, Bertolotti A. Coping with Protein Quality Control Failure. Annu Rev Cell Dev Biol. 2017 10; 33:439–465. doi:10.1146/annurev-cellbio-111315-125334.

Pilon M, Schekman R, Römisch K. Sec61p mediates export of a misfolded secretory protein from the endoplasmic reticulum to the cytosol for degradation. EMBO J. 1997 Aug; 16(15):4540–8. doi:10.1093/emboj/16.15.4540.

Ritchie ME, Phipson B, Wu D, Hu Y, Law CW, Shi W, Smyth GK. limma powers differential expression analyses for RNA-sequencing and microarray studies. Nucleic Acids Res. 2015 Apr; 43(7):e47. doi:10.1093/nar/gkv007.

Römisch K. Endoplasmic reticulum-associated degradation. Annu Rev Cell Dev Biol. 2005; 21:435–56. doi:10.1146/annurev.cellbio.21.012704.133250.

Römisch K. A Case for Sec61 Channel Involvement in ERAD. Trends Biochem Sci. 2017 03; 42(3):171–179. doi:10.1016/j.tibs.2016.10.005.

Savitski MM, Reinhard FBM, Franken H, Werner T, Savitski MF, Eberhard D, Martinez Molina D, Jafari R, Dovega RB, Klaeger S, Kuster B, Nordlund P, Bantscheff M, Drewes G. Tracking cancer drugs in living cells by thermal profiling of the proteome. Science. 2014 Oct; 346(6205):1255784. doi:10.1126/science.1255784.

Schäfer A, Wolf DH. Sec61p is part of the endoplasmic reticulum-associated degradation machinery. EMBO J. 2009 Oct; 28(19):2874–84. doi:10.1038/emboj.2009.231.

Schäuble N, Lang S, Jung M, Cappel S, Schorr S, Ulucan Ö, Linxweiler J, Dudek J, Blum R, Helms V, Paton AW, Paton JC, Cavalié A, Zimmermann R. BiP-mediated closing of the Sec61 channel limits Ca2+ leakage from the ER. EMBO J. 2012 Aug; 31(15):3282–96. doi:10.1038/emboj.2012.189.

Scheper W, Thaminy S, Kais S, Stagljar I, Römisch K. Coordination of N-glycosylation and protein translocation across the endoplasmic reticulum membrane by Sss1 protein. J Biol Chem. 2003 Sep; 278(39):37998–8003. doi:10.1074/jbc.M300176200.

Schwanhäusser B, Busse D, Li N, Dittmar G, Schuchhardt J, Wolf J, Chen W, Selbach M. Global quantification of mammalian gene expression control. Nature. 2011 May; 473(7347):337–42. doi:10.1038/nature10098.

Servas C, Römisch K. The Sec63p J-domain is required for ERAD of soluble proteins in yeast. PloS One. 2013; 8(12):e82058. doi:10.1371/journal.pone.0082058.

Shamu CE, Walter P. Oligomerization and phosphorylation of the Ire1p kinase during intracellular signaling from the endoplasmic reticulum to the nucleus. EMBO J. 1996 Jun; 15(12):3028–39.

Sikorski RS, Hieter P. A system of shuttle vectors and yeast host strains designed for effcient manipulation of DNA in Saccharomyces cerevisiae. Genetics. 1989 May; 122(1):19–27.

Stein A, Ruggiano A, Carvalho P, Rapoport TA. Key steps in ERAD of luminal ER proteins reconstituted with purified components. Cell. 2014 Sep; 158(6):1375–1388. doi:10.1016/j.cell.2014.07.050.

Stirling CJ, Rothblatt J, Hosobuchi M, Deshaies R, Schekman R. Protein translocation mutants defective in the insertion of integral membrane proteins into the endoplasmic reticulum. Mol Biol Cell. 1992 Feb; 3(2):129–42. doi:10.1091/mbc.3.2.129.

Tretter T, Pereira FP, Ulucan O, Helms V, Allan S, Kalies KU, Römisch K. ERAD and protein import defects in a sec61 mutant lacking ER-lumenal loop 7. BMC Cell Biol. 2013 Dec; 14:56. doi:10.1186/1471-2121-14-56.

Vashist S, Ng DTW. Misfolded proteins are sorted by a sequential checkpoint mechanism of ER quality control. J Cell Biol. 2004 Apr; 165(1):41–52. doi:10.1083/jcb.200309132.

Vembar SS, Brodsky JL. One step at a time: endoplasmic reticulum-associated degradation. Nat Rev Mol Cell Biol. 2008 Dec; 9(12):944–57. doi:10.1038/nrm2546.

Voorhees RM, Hegde RS. Structure of the Sec61 channel opened by a signal sequence. Science. 2016 Jan; 351(6268):88–91. doi:10.1126/science.aad4992.

Walter P, Ibrahimi I, Blobel G. Translocation of proteins across the endoplasmic reticulum. I. Signal recognition protein (SRP) binds to in-vitro-assembled polysomes synthesizing secretory protein. J Cell Biol. 1981 Nov; 91(2 Pt 1):545–50.

Wang L, Dobberstein B. Oligomeric complexes involved in translocation of proteins across the membrane of the endoplasmic reticulum. FEBS Lett. 1999 Sep; 457(3):316–22.

